# Comparative efficacy of antiviral strategies targeting different stages of the viral life cycle: A viral quasispecies dynamics study

**DOI:** 10.1101/2022.10.10.511620

**Authors:** Pancy Lwin, Greyson R. Lewis, Moumita Das, Barbara A. Jones

## Abstract

While the COVID-19 pandemic continues to impact public health worldwide significantly, the use of antiviral drugs and therapies has dramatically reduced the instances of severe disease and death. More broadly, the unprecedented use of antivirals also provides hope for preventing and mitigating similar viral outbreaks in the future. Here we ask: What are the comparative impact of antiviral therapeutics targeting different stages of the viral lifecycle? How do antiviral therapeutics impact the viral population in the bloodstream, or in other words, the viral load in high and low-immunity individuals? To address these questions, we use a model of viral quasispecies dynamics to examine the efficacy of antiviral strategies targeting three critical aspects of the viral life cycle, fecundity, reproduction rate, or infection rate. We find a linear relationship of the viral load with the change in fecundity and a power law with the change in the reproduction rate of the virus, with the viral load decreasing as the fecundity and the reproduction rates are decreased. Interestingly, however, for antivirals that target the infection rate, the viral load changes non-monotonically with the change in infection rate; the viral population initially increases and then decreases as the infection rate is decreased. The initial increase is especially pronounced for individuals with low immunity. By examining the viral population inside cells for such cases, we found that the therapeutics are only effective in such individuals if they stop the infection process entirely. Otherwise, the viral population inside cells does not go extinct. Our results predict the effectiveness of different antiviral strategies for COVID-19 and similar viral diseases and provide insights into the susceptibility of individuals with low immunity to effects like long covid.

## Introduction

In the evolution of viruses, especially in error-prone RNA viruses, the interplay of mutations, replications, and selection processes lead to populations of viruses, known as a viral quasispecies, with similar sequences but a distribution of genomes [1]. Statistical mechanics of viral quasispecies provides a framework to understand how a virus survives the immune clearance system and provides insights into the population biology of viruses. In addition, quasispecies dynamics can suggest approaches for developing antivirals because the viruses can be considered “moving targets” when replicating [2]. In this work, we use this framework to systematically study the impact of different anti-viral therapeutic strategies that target different aspects of the life-cycle of respiratory viruses such as influenza, cold viruses and particularly SARS-CoV-2. We build on previous studies [3],[4] that model the viral life cycle as three main discrete stages of infection, immune clearance and reproduction, and investigate the aftermath of administering antivirals at a specific time after first infection, that selectively reduce the infection rate, the reproduction rate, or the fecundity of the virus, where we define fecundity as the maximum number of new viruses that can be produced from each progenitor virus. While we are investigating a general rule for a broad class of viruses, the results provide useful insights on which part of the viral cycle to focus on to produce the most effective antiviral therapeutics.

Antivirals work by disrupting specific aspects of the viral life-cycle and can be generally categorized based on their strategy [5]. Inhibitors of viral RNA polymerase or RNA Synthesis may be able to reduce the reproduction rate or fecundity of viruses [6]. Therapeutics can inhibit viral protein synthesis such as processes via which polypeptide chains are broken down into viral proteins by proteases, which is an essential step in viral replication. The inhibitors bind to the protease enzyme and disrupt their functioning preventing viral assembly [7]. As a result, it prevents the formation of new viruses. These antivirals can affect the reproduction rate of the viruses. Examples include CRISPR-Case-based Gene Therapy [8], defective viral genomes (DVGs) [9], monoclonal antibodies and other protease inhibiting antivirals [10, 11, 12, 13, 14, 15]. Some antivirals may restrict the entry of viral protein into the nucleus thereby reducing the reproduction of viruses [16]. In addition, some viral entry inhibitors may restrict viral entry via membrane fusion or endocytosis [7], [17], [18]. These antivirals may be able to change the probability of infection. Note that the above-mentioned antivirals are not necessarily proven effective against Covid-19 patients. Here we consider them as examples of different types of antivirals affecting fecundity, reproduction and infection rates, as shown in Figure 1.

**Figure 1:**
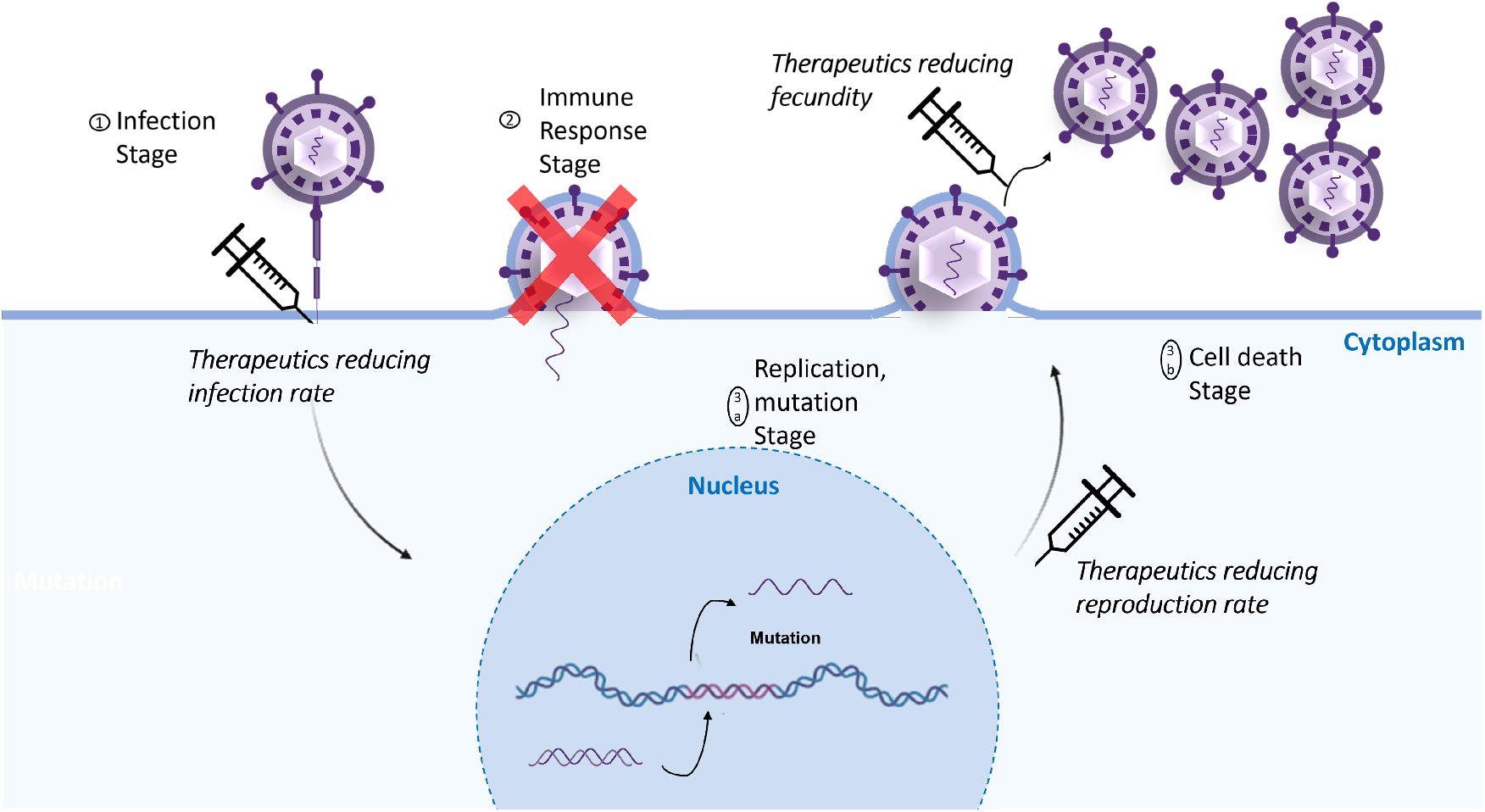
Virus Life Cycle and Different Therapeutics in Different Stages of the Cycle. The viral life cycle is described in a simplified schematic, including the viral therapeutics working on the cycle. The cell occupation probability and the virus population are calculated during **1) Infection, 2) Immune Clearance** and **3) Reproduction**. We modeled the actions of the therapeutics as effects on the infection rate, fecundity and reproduction rate.

There have been numerous modeling efforts focused on understanding and incorporating different facets of the viral infection along with the therapeutics, and we summarize some key classes of them here. Early models centered around within-host viral kinetics (VKs), which accounts for the time evolution of uninfected and infected cells with virions [19, 20]. This model was initially used in studying HIV-1 infections [21] and was later extended to other viral infections. Moreover, PK/VK models incorporating Pharmacokinetics (PK) dynamics and Hill coefficient have been developed to study the antiviral effectiveness [22, 23, 24]. Applications of such models include the assessment of alisporivir interferon-free treatment in hepatitis C virus infected patients [25]. Similar models are also being used in studying Covid-19 disease dynamics [26, 27, 28]. In addition to the mechanistic approaches mentioned above, more data-driven methods of learning about SARS-CoV-2 variant binding strengths are being carried out through simulations [29]. Lastly, for Model Informed Drug Development (MIDD), some other modeling efforts [30, 31] involve using Quantitative System Pharmacology principles (QSP).

In our model, we consider an iterative procedure, shown in Figure 1, in which a virus first attempts to infect a cell from an environment composed of a distribution of viral types. We discuss more about this distribution, a quasi-species distribution, below. If the virus is successful in infecting the cell, it is next exposed to the host immune system. Any viruses surviving that then have a probability of reproducing with a fecundity, with each offspring virus given a random mutation on one codon. These viruses then become the new environment, and go on to try to infect other cells.

In our model system, the cells are all identical, with a “lock” 50 codons (AA) long. Each codon takes a value from A-Z, for the cells a fixed pattern with a 50-character catch phrase. The viruses, on the other hand, have a “key” 100 codons long, again with each taking the 26 values A-Z, but for the viruses the values are variable, allowing 26^100^ different possible viruses. In the infection process in our model, the virus aligns its key with the cell’s lock, sliding its 100 codons along the lock and testing at each alignment how many codon matches there are. The alignment with the greatest number of matches of virus to cell has a number of matches we term the “match number” for that virus, and is the identifier for that virus. Because of the relative sizes of codon regions between virus and cell, the match number can range from 0 to 50, giving 51 different values.

As the viruses enter from the environment, they hop from cell to cell, at each trying to infect. If a cell is already full, the virus cannot infect and must hop to the next one. If after trying all cells the virus cannot infect, it is considered to leave the active part of the host. The probability of a virus entering a cell on a given hop, if the cell is not full, is ***e***_***m***_, where 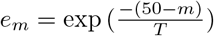. Here *m* is the match number for the virus attempting to infect, and T is a parameter we term cell permissivity, a measure of how easily a virus with fixed match number can enter a cell. For low permissivity, the match numbers need to be near 50 to infect, while for high permissivity, a broad range of viruses are able to infect. The net result of the hopping and probabalistic infection of N successive viruses from the environment, with the fixed number of cells gradually filling up as the later viruses try to infect, is an updated infection probability of the cells, and a new distribution of viral match numbers in the cells, which we term *ψ*^*I*^ (*m*). This is expressed in a recursive formula which is described in the Supplementary Information (SI).

The second stage or the energy barrier to overcome for the virus after the infection is the immune response. It is modeled as a sigmoid, which will generate a high immune response for any virus with more than a few matches, 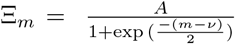. The immunity parameter A is varied between 0 and 1, while the onset parameter *?* has been suggested to be set to 6, as a typical epitope length [1]. Therefore, Ξ_*m*_ will approach 1 when m is a moderate number, and A is 1. The cell occupancy after the immune response is calculated using the above-mentioned sigmoid function, *ψ*^Ξ^(*m*) = *ψ*^*I*^ (*m*)(1 – Ξ_*m*_). While infection increased the number of viruses in the cells, the immune response decreases the number of viruses in the cells, and again changes the distribution of *m* in the cells. Note that infection and immune response create opposite pressures on the virus, one for higher match number to infect, but then needing low match number to escape the immune process.

The final stage is the reproduction stage, where the viruses in the cells have a probability based on their match number to reproduce and mutate. We take the probability to reproduce to be ***e***_***m***_. Then the viruses left in the cells can be expressed as *ψ*^*R*^(*m*) = *ψ*^Ξ^(*m*)(1 − *e*_*m*_). Conversely, the probability of the virus producing an offspring successfully is *ψ*^*F*^ (*m*) = *e*_*m*_*ψ*^Ξ^(*m*). We set the fecundity to be 20, and for each viral offspring, exactly one of the 100 codons is randomly changed. This change can result in the virus having a new match number which is either one greater than before, one less than before, or unchanged. This three-way probability is different for every m (more likely to increase for small m, more likely to decrease for larger m), and the net we express as a mutation matrix applied to each probability vector *ψ*^*F*^ (*m*). This cohort of reproduced viruses then composes the viral environment for the next iteration. The number of viruses in the environment for each *m* is *cfψ*^*F*^ (*m*), where *c* is the number of cells and *f* is the fecundity. Each iteration is taken to be one time step. Iterations are repeated until steady state is achieved. By varying immune strength *A* and cell permissivity *T* over a full range, a phase diagram can be obtained.

We use this framework discussed above to examine three strategies for anti-viral therapeutics: reduction of infective ability, reproductive ability, and fecundity. To reduce the reproduction rate and fecundity, we multiply the previously defined *ψ*^*F*^ (*m*) and *f* by scaling prefactors *R* and *F* respectively, with *R* and *F* ranging from 1 (no therapeutic) to zero. To reduce the infection rate, we multiply the part of the infection process that has to do with the ability to enter the cell by a scaling prefactor I ranging between 1 and 0. Details of how this is implemented are in the Supplementary Information. Every simulation starts at time *t* = 0 with the parameters *I, R*, and *F* set to unity, and then at the point *t* = 10 one or the other of the three anti-viral parameters is set to a value less than one and kept fixed at this value, indicating the continual dosing of a therapeutic. The simulation is continued until a steady state is reached, in which either the virus is extinct (successful therapeutic) or a steady state nonzero level of virus remains. We vary the reduction parameters *I, R*, and *F* over a range of values between 0 and 1, and we choose four characteristic places on the phase diagram (sets of parameters *A* and *T*) to have a sampling of hosts with both low and high immunity, and low and higher cell permissivity.

The full phase space of the model discussed in this work has three main phases of disease: acute, opportunistic, and chronic [4]. Here we focus on the acute phase representing respiratory diseases including flu, SARS, and in particular COVID-like diseases. Within this phase, we sample from four different locations as shown in Figure 2(c). The corresponding four sets of permissivity (*T*) and immunity (*A*) used are (*T* = 0.5, *A* = 0.2), (*T* = 0.5, *A* = 0.5), (*T* = 0.5, *A* = 8) and (*T* = 12.9, *A* = 0.5). We choose three locations at the same permissivity of *T* = 0.5 which is in the middle of the phase. The last point we choose is at the permissivity of *T* = 12.9 and immunity of *A* = 0.5.

**Figure 2:**
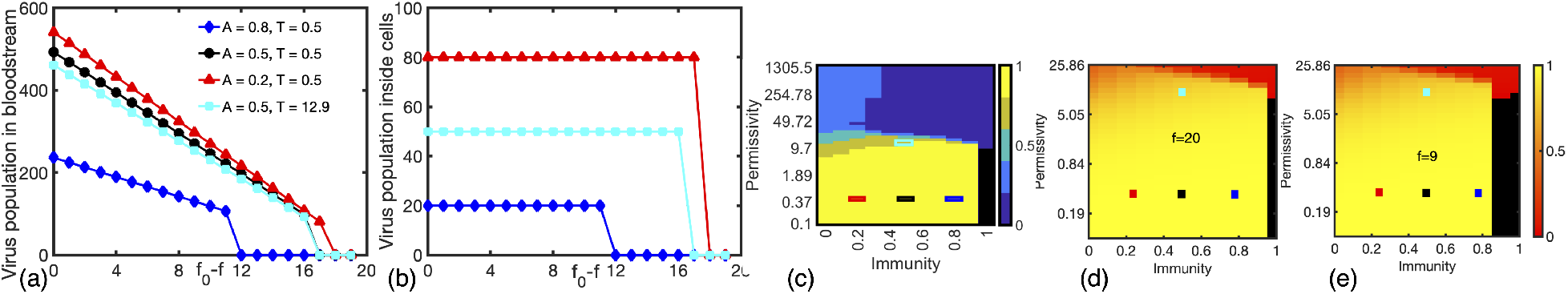
Relationship between virus populations and the change in fecundity *f*. (a) Virus population in the bloodstream and (b) virus population inside cells at steady state as a function of the change in fecundity, *f*_0_ − *f* following the application of the antiviral. (c) Heat map showing distinct regions for acute, chronic and opportunistic phases as described in [4]. The four rectangular boxes in red, black, blue and cyan show the four cases studied in this paper, which focus only on the acute phase (yellow in (b)). (d) Heat map of the order parameter for a control sample where no antiviral is used and (e) for the case where the antiviral is administered; both heat maps have the same color-scale. The order parameter is the normalized mean of the match number. Extinct viruses are marked with a black color on the heat map.

## Results

First, we discuss the effect of lowering fecundity through therapeutics as shown in Fig. 2. The viral population in the bloodstream, also called the viral load, decreases linearly with the decrease in fecundity, *f*_0_ − *f*, in all cases studied (Fig. 2(a)). The viral population in the cells is initially much smaller than in the bloodstream and stays almost steadily constant as the fecundity is reduced; however, it goes to zero (extinct virus) at the same level of therapeutic as that for the viral population in the bloodstream ((Fig. 2(b)). The viral population both in the bloodstream and inside cells is the smallest for the case where the immunity is highest and the permissivity is lowest (*A* = 0.8, *T* = 0.5), and it reaches zero for a smaller reduction in fecundity for this case, compared to all other cases. For a fixed permissivity (*T* = 0.5), the viral populations increase with decreasing immunity, *A*, as expected. For fixed immunity (*A* = 0.5), the dependence on permissivity, *T*, is does not show a clear, monotonic trend which is shown and discussed in the Supplementary Information. We also calculated the order parameter in the parameter space of immunity and permissivity for high (*f* = 20) and low (*f* = 9) fecundity. We find that the parameter space over which the virus goes extinct grows as the fecundity is reduced as shown in heat map in Figs. 2(d) and 2(e). However, the specific regions of phase space we discussed above do not show a significant change in order parameter.

To understand the impact of antivirals which lower the probability of reproduction of viruses, we changed the reproduction rate of the system, *R*, after 10 time steps. At the default value of *R* = 1, i.e., before the anti-viral is administered, the case with the lowest A and the lower T generates the most viral load in the environment. As with fecundity, we observed a reduction in the viral population in the bloodstream as we decrease the reproduction rate, *R*, for all cases (Fig. 3(a)), while the viral population in cells initially stays constant as *R* is decreased but jumps to zero at the same time as the viral population in the bloodstream (Fig. 3(b)). The decrease in the viral population in the bloodstream, however, follows a non-linear relationship with *R*_0_ − *R*, unlike with decreasing fecundity (Fig. 3a). This non-linear scaling is the most noticeable for the case of *A* = 0.2 and *T* = 0.5, when there is a competition between low immunity and low permissivity. The order parameter heatmap for the reduced reproduction rates show an increase in the extinction region as in case of reducing fecundity (Fig. 3(c)).

**Figure 3:**
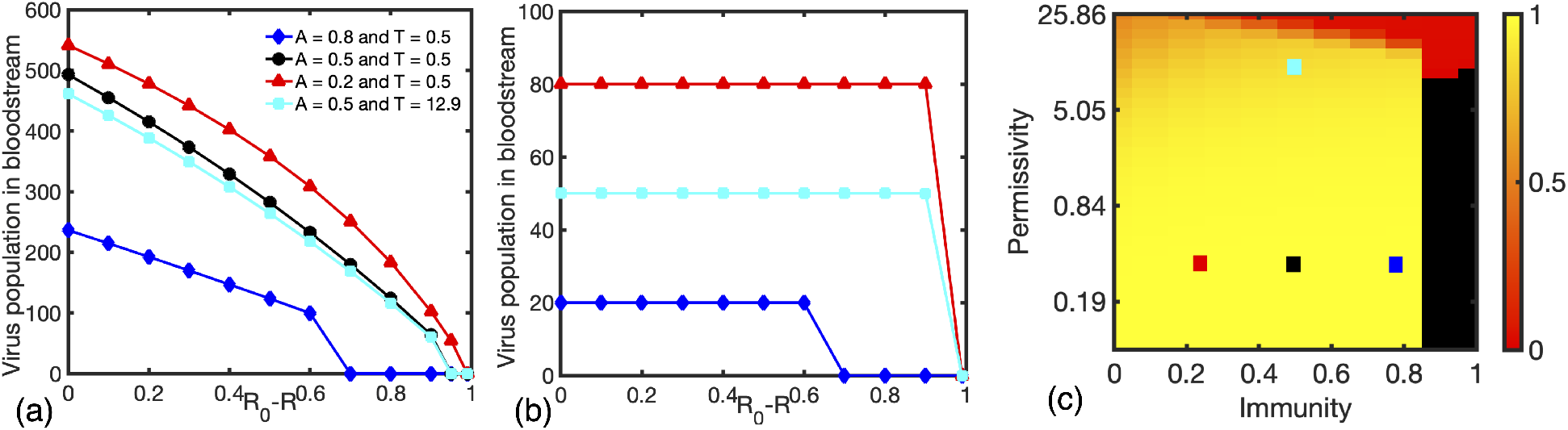
Relationship between virus populations and the change in Reproduction Rate *R*. **(a)** The virus population in the bloodstream and (b) in the cell at steady state as a function of the change in the reproductive rate, *R*_0_ − *R*. (c) The heat map of the order parameter shows an increase in the area of extinction when the viral reproduction rate is reduced. Extinct viruses are marked with a black color on the heat map. All heat maps have the same color-scale.

We investigated the impact of antivirals which target the rate of infection by implementing a reduction in the infection rate at time= 10 in our simulations. The resulting impact was unexpected and different from the antivirals that had targeted fecundity and reproduction rate. Here, the viral population in the bloodstream changes non-monotonically as we decrease the infection rate, first increasing and reaching a maximum which can be quite marked, and then decreasing and finally going to zero i.e. going extinct, as shown in Fig. 4(a). We conjecture this initial increase takes place because even though a decrease in infection rate leads to fewer viruses in cells, these viruses now face less competition and able to reproduce and fight the immune system in the absence of any reduction in those processes, effectively leading to an initial increase in the viral load. This initial increase in the viral load is largest for the case of the lowest immunity *A* = 0.2. The change in the viral population inside cells follows qualitatively similar trends in this case as with the reduction in fecundity and reproduction rate (Fig. 4(b)). For both viral populations in the bloodstream and in cells, for higher immunity, viral extinction requires smaller reductions in infection rate (Fig. 4(a) and (b)). This is also observed in the heat map of the ordered parameter, for the cases *I* = 0.1050, *I* = 0.0235, and *I* = 0.0052 shown in Fig. 4 (c),(d), and (e)).

**Figure 4:**
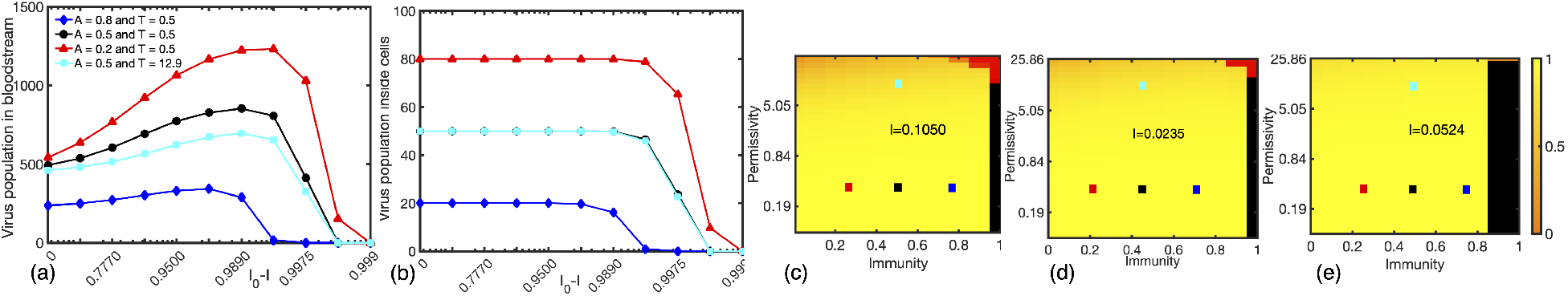
Relationship between virus populations and the change in Infection Rate *I*. **(a)** The virus population in the bloodstream and **(b)** in the cell at steady state versus the reproductive rate *I* changed to after the tenth time step. The antiviral reducing infection rate shows significantly different trends in all cases. **(c)**,**(d)**, and **(e)** Phase diagrams with various levels of therapeutic. Extinction regions are marked with a black color on the heat map. The chosen infection rate is indicated on each heat map.

A comprehensive analysis of the impact of the therapeutics in terms of the evolution of the viral population, match number, and the quasi-species distribution is shown in Fig 5. For antivirals that reduce the fecundity and reproduction rate, for the case of high immunity, the viral load drops to a value close to zero after the antiviral is administered and stays there, while for the case of low immunity although the viral load drops right after the antiviral is used, it does not go to zero, and reaches a non-extinct steady state. No clear mutations in the quasispecies distribution are observed. Our results for the time dependence of viral load for fixed values of infection-reducing therapeutic chosen at two characteristic values in the rise and fall show that the viral load increases quite quickly after the therapeutic is implemented for high immunity, but show an extended rise over time for the lower immunity. The only time this trend is reversed and we see a decrease is for the case that combines high immunity (*A* = 0.8) with low permissivity (*T* = 0.5) and low infection rate (*I* = 0.00524) (Fig. 5 (m)). A possible explanation for the rise in viral load with implementation of infection-lowering therapeutic is suggested by the studies of quasi-species distribution as a function of time, showing a set of mutations most prominently seen in (l) and (p) to higher match number, mutations that allow the virus to reproduce more. Although the mutations do not appear particularly large, a reminder of the *e*_*m*_ function governing reproduction, which is an exponential function of the deviation of the viral match number from *m* = 50. As a result, we see the virus population increasing. We provide more details in the supplementary information.

**Figure 5:**
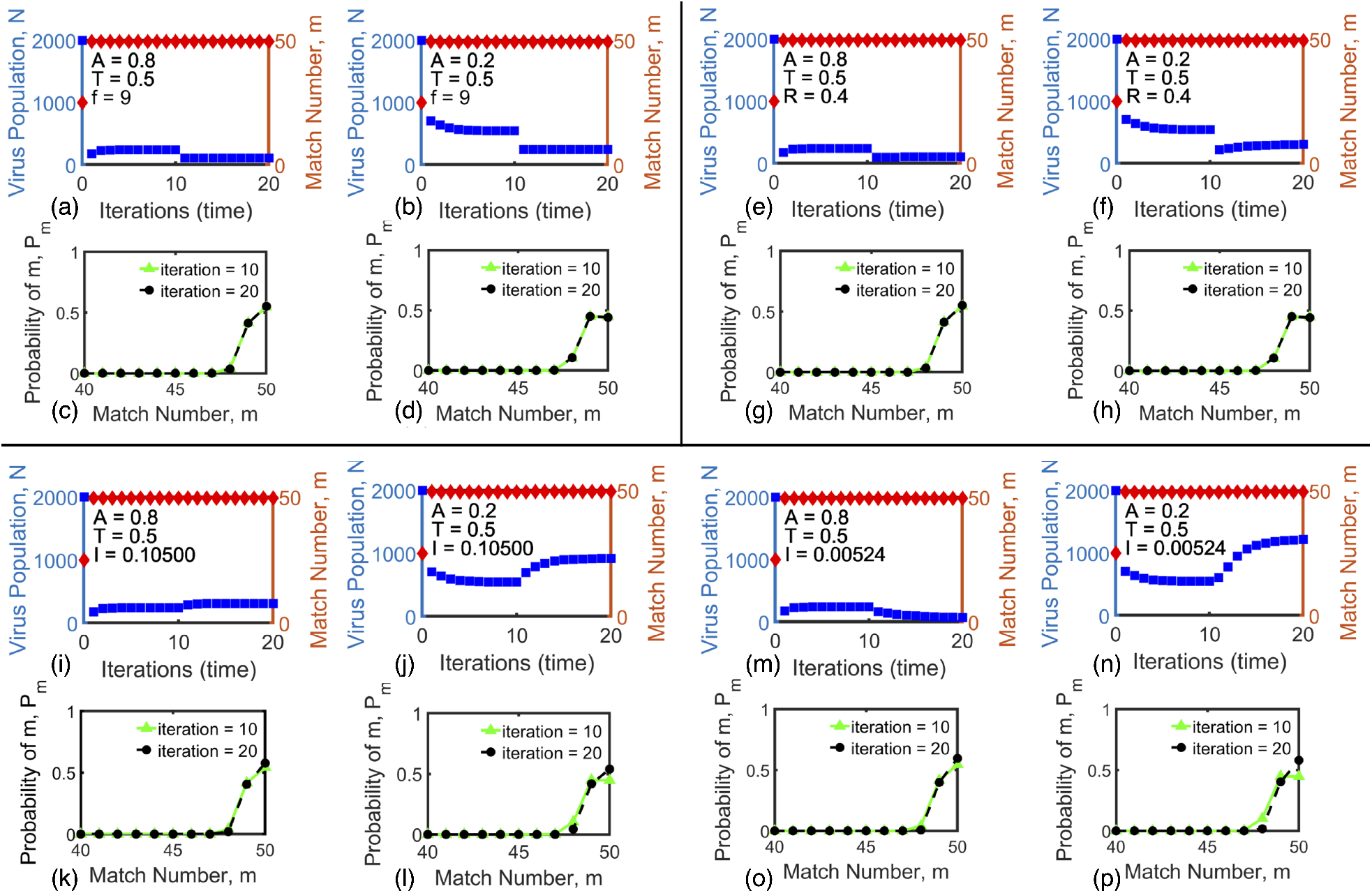
The virus population and the probability of matches for given permissivity and immunity for antivirals targeting the fecundity, reproduction rate, and infection rate. Figures (a)-(d) show the impact of varying the fecundity (lowering it to *f* = 9), figures (e)-(h) the results of varying the reproduction rate (lowering it to *R* = 0.4), and figures (i)-(p) show the results of varying the infection rate (lowering it to *I* = 0.10500 for (i)-(l), and to *I* = 0.00524 for Fig. (m)-(p)). The virus population in the bloodstream (viral load) and average match number (order parameter) are shown in Figs (a),(b),(e),(f),(i),(j), (m), and (n), and the quasispecies distributions before and after the therapeutic is administered are shown in Figs. (c), (d), (g), (h), (k),(l)(o),(p). The permissivity is kept fixed in all cases at *T* = 0.5. Figs (a),(c),(i),(k),(e),(g),(m),(o) show results at a high value of immunity *A* = 0.8, while Figs. (b),(d),(j),(l), (f),(h),(n),(p) show results at a low value of immunity *A* = 0.2. Note that no clear mutations in the quasispecies distribution are observed for the cases of lowering fecundity or reproduction rate, however for antivirals that lower the infection rate, mutations in the quasi-species distribution are observed with a shift to higher match numbers enabling higher reproduction rate, particularly for lower immunity.

Finally, we examined how the viral population in the bloodstream and in cells changed over time in response to antiviral therapeutics that lowered the fecundity, the reproduction rate, and the infection rate, for the cases of high and low immunity. In Fig 6, *F* = 0.05, *R* = 0.01 and *I* = 0.0015 are selected to show how the virus population in both bloodstream and cells approach extinction. As seen in the first column, with the high immunity *A* = 0.8, the viral load reduces abruptly; therapeutics and high immunity work together to eliminate viruses. However, with the low immunity *A* = 0.2, as shown in the second column, viruses stay inside the cell reservoir for dozens of our time units (many days, weeks even) before viral population approaches a tiny number, if it does. (As shown for the therapeutic affecting infection, even for a reduction of ability to enter a cell to 0.1 percent of its full value, the virus does not go extinct, and in fact rises with time up to its steady state value. In the SI we show the reduction of infection to 0.00524 still does not clear the virus for the low-immunity sample.) We postulate that the time delay of viral extinction in the bloodstream and inside cells might give insights into the initiation of long Covid and provide valuable contributions to discussion about the nature of illness.

**Figure 6:**
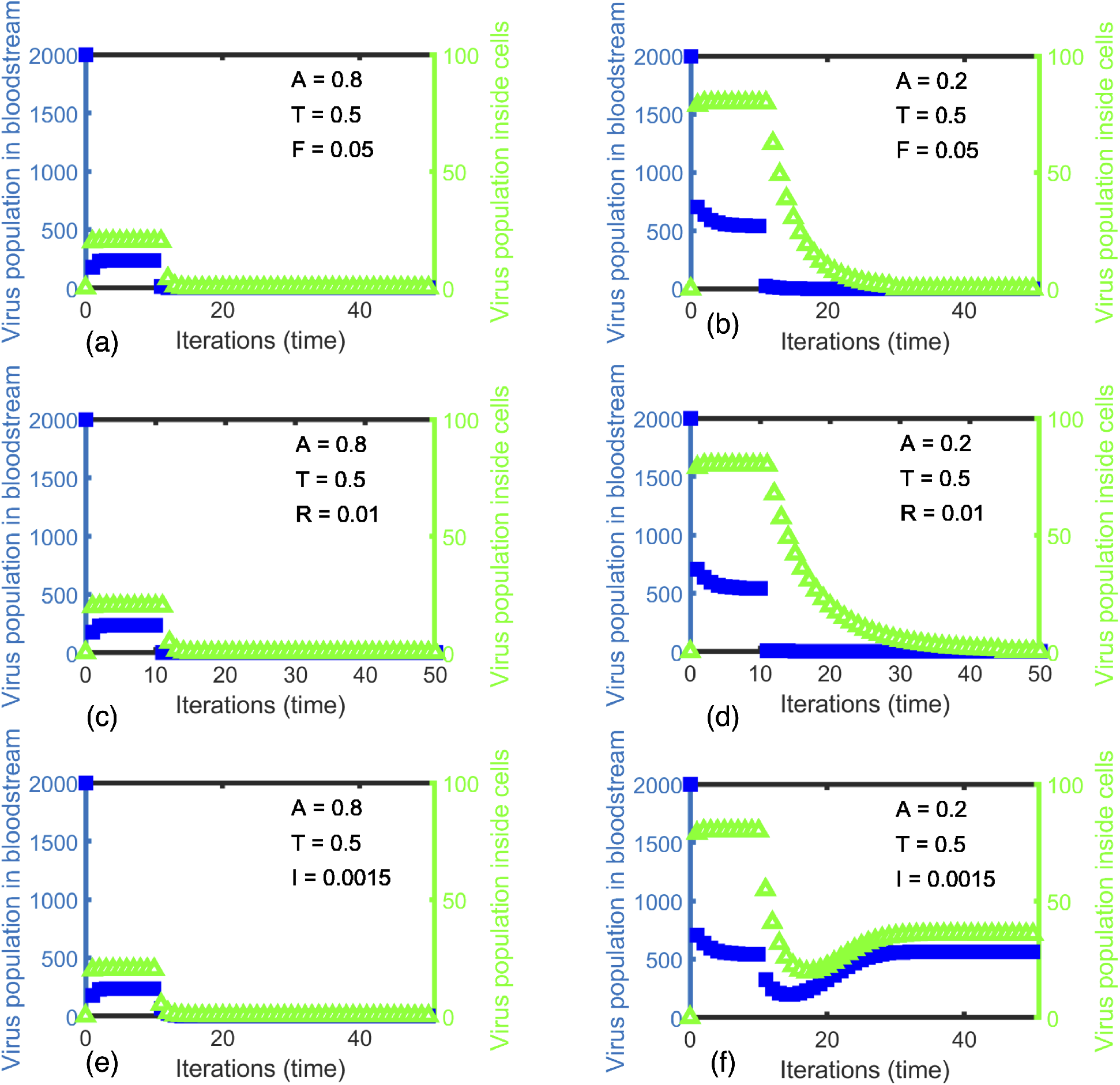
Time-dependent comparison of virus population in bloodstream vs inside cells at extinction. The virus population in the bloodstream plotted in blue (right y-axis) and inside cells plotted in green (left y-axis). Permissivity *T* = 0.5 in all cases. Immunity *A* = 0.8 for the left columns (a), (c), (e) and Immunity *A* = 0.2 for the right columns (b), (d), (f)

## Discussion

We investigated the efficacy of antivirals in terms of their ability to reduce the fecundity, the reproduction rate and the infection rate of viruses by implementing the change in the respective parameters after a certain number of iterations, representing a time delay from first infection to feeling sick enough to seek medical intervention. As expected, an antiviral that leads to a fewer number of offspring per virus, i.e. a smaller fecundity, leads to a decrease in the viral load. This effect is monotonic, and the number of viruses at a given time is a linear function of the change in fecundity; this is a desired behavior because it allows to predict the impact of the antiviral with consistency, including to infer the data in the regions parameter space we do not show in the manuscript. For the therapeutics reducing the reproduction rates of the viruses, once again the viral load decreases smoothly monotonically with the decrease in the reproduction rate, but this decrease is linear only for the case of high immunity and low permissitivity (*A* = 0.8, *T* = 0.5). An example of an antiviral that reduces the reproduction rate is the the FDA approved Paxlovid, for mild and moderate symptom of Covid-19 patients [32]. It combines, nirmatrelvir, a protease inhibitor which interferes the replication process, and ritonavir, which makes sure nirmatrelvir is not metabolized away easily [33]. The success of this antiviral in preventing progression of the disease and reducing hospitalization agrees with our results on the effectiveness of antiviral strategies that target the reproduction rate of viruses.

The impact of antivirals that lower the ability to infect was intriguing and somewhat counterintuitive. Here the viral load changed non-monotonically with the decrease in infection rate, initially increasing, reaching a maximum, and then decreasing. We conjecture this initial increase to be due to less competition faced by the viruses already successful in infecting the cell, their unrestricted reproduction, and potential mutations; the latter is suggested by the shift to increased match number of the quasi-specicies distribution when the therapeutic is administered. These effects combine leading to an accumulation of viruses in cells, with the viruses reproducing until all the cells are empty and devoid of viruses, which critically depends on the level of immunity. An example of an antiviral with a similar strategy is Ivermection [34], which despite its reported ability to reduce the infection rate of a wide range of viruses, was found to be not be very effective in mitigating Covid -19 in patients. This is likely because, as our results highlighted, reducing the infection rate is not effective in reducing the overall viral load if the viruses continue to replicate, and potentially mutate.

One question left is what defines and quantifies the illness or symptoms of the viral infection. If the viral load is reduced in the bloodstream but most cells are still filled with viruses, there is a possibility that the patient may still suffer from the symptoms of the viral infection. For Covid-19, many patients report the long haul struggle with the symptoms such as trouble breathing, muscle aches and chronic fatigue [35]. Despite the fact that the causes can be varied from blood clots [36, 37] to immune abnormalities [38], some research suggest that antivirals would be a solution to eliminate the reservoir of viruses inside cells [39]. Our model also shows the slow approach of the viruses inside the cells to extinction under therapeutics. The extinction rate depends on the strength of the applied therapeutics and very much on the immune strength. Individuals in our model with high immunity not only get well, but get well rapidly after therapeutic is administered (See Fig. 6 a,c and e). For individuals in our model with a weak immune response, both in Fig. 6 b and d, reducing the fecundity and reproduction rate to a very small number, which is equivalent to administering a strong therapeutic, brings the virus population in the bloodstream within a few iterations to extinction. However, there is a much longer time scale to clearing the virus from the cells for those with low immunity, extending to 20-30 iterations (approx. 10-30 days). And as Fig. 6 f shows, for some therapeutics, even a strong dose for those with low immune response clears the virus from neither blood stream nor cells. In summary, those with weaker immunity not only don’t always get their virus cleared if the therapeutic is not strong enough, but also, most tellingly, have a very long time scale to the removal of virus from their cells. Since it is viruses in cells that activates the immunity, our results may give insights to both exaggerated immune response such as a cytokine storm, and as well as the effects of long Covid.

### Methods

The simulations are performed on Rochester Institute of Technology’s Hight Performance Computing (HPC) cluster, named SPORC (Scheduled Processing On Research Computing). With the help SLURM manager and the capacity of the cluster, each task for each simulation of permissivity T and immunity A combination is run efficiently. The system size is defined by maximally 2000 number of viruses attempting to infect 100 empty cells for 400 iterations. The convergence is shown by reaching the steady state values in the number of viruses in the environment.

## Acknowledgements

PTL was partially supported by the National Science Foundation award CBET-1604712, and an INTERN supplement to this award.

