## Supplementary Information for "Comparative efficacy of antiviral strategies targeting different stages of the viral life cycle: A viral quasispecies dynamics study"

October 10, 2022

#### 1 Mathematical model

We explain the details of the viral life cycle models we used in our main paper. There are three different stages of a viral life cycle we simulated - infection, immune clearance and reproduction. We calculate the probability distribution of viral quasi-species in each stage with or without the applied therapeutics.

##### 1.1 Details of infection stage

The probability distribution of match numbers in the cells after infection by a quasi-species distribution of viruses is calculated as follows. In our model the virus quasispecies distribution infects via viruses emerging from the blood stream, and attempting to infect cells, hopping across cells, trying to infect one cell at a time. At each cell, there are two constraints: the first is related to the cell permissivity, represented by $e_{m(infect)}$, and second to whether the cell is already infected and occupied and this is represented by the sum over all m of the quasispecies present in the cell prior to virus $k$ arriving, or $\sum_{m'} \psi^I_{k-1}(m')$ . Both of these quantities are between 0 and 1, representing probabilities. The probability that the virus will find the cell empty enough to enter is then $1 - \sum_{m'} \psi^I_{k-1}(m')$, and the combined probability to infect a cell in a single attempt is the product of the two, $e_{m(infect)}(1 - \sum_{m'} \psi^I_{k-1}(m'))$.

The probability of the virus encountering all $c$ cells and infecting at most one cell is expressed as one minus the probability of encountering all $c$ cells and not infecting any one of them, or $[1 - [1 - e_{m(infect)}(1 - \sum_{m'} \psi^I_{k-1}(m'))]^c]$. We average over all cells and the full viral quasispecies distribution. The net distribution in the cells after the $k^{th}$ virus has tried to infect is then the distribution after the encounter with the $(k-1)^{th}$ virus plus the new infection attempt from virus $k$. This distribution is shown in the following equation:

$$\psi^I_k(m) = \frac{P_m}{c}[1 - [1 - e_{m(infect)}(1 - \sum_{m'} \psi^I_{k-1}(m'))]^c] + \psi^I_{k-1}(m)$$

This process repeats until all $N$ viruses have tried to infect, so that the final $\psi^I(m) = \psi^I_N(m)$. The process starts with $\psi^I_{k=0}(m) = \psi^R(m)$, the viruses remaining in the cells after the previous iteration (previous time step), and the first virus is represented by $k = 1$.

This process assumes an integer number of viruses $N$, but the results of the previous iteration (reproduction and mutation process) yield a real number. To obtain an integer number of viruses we interpolate to the nearest integer and define this as N.

#### 1.2 Administering Anti-viral Therapeutics

For viral fecundity and reproduction, there is only one parameter affecting the probability of viral success (in our model), and we multiply the respective probability prior to administering the antiviral by a prefactor representing dosage of the therapeutic, $F$ and $R$. For therapeutics affecting infection, of the two factors that impact viral infection in our model, cell permissivity and cell occupancy, only cell permissivity is modified. Therefore, we do not multiply the entire infective addition of each $k$th virus by the inhibition of the therapeutic $I$, but rather just that part involved in cell permissivity, $e_m = exp(\frac{-(50-m)}{T})$. We write $e_{m(infect)}$ in the equation for infection above, to show where it appears. The equations representing the application of the therapeutics for each of the three strategies considered in this paper are shown below.

$$\begin{aligned} e_{m(infect)} &= I\ e_m \\ e_{m(repro)} &= R\ e_m \\ f_{therapeutic} &= F\ f \\ N &= F\ cf \sum_m \psi^F(m) \end{aligned}$$

Here $I$, $R$, and $F$ are all between 1 (no therapeutic) and 0. The last equation shows how the number of viruses, $N$, in the bloodstream for the next iteration is obtained, where $f$ is the fecundity. Note that $N$ above is the expectation value of the number of viruses, and hence a real number. This is rounded to the nearest integer for the infection process, where $k$ must run from 1 to $N$.

#### 1.3 Meaning of extinction and time in the model

Here we are going to elaborate the definition of extinction values and time. In our model, we assumed that the virus is extinct when the virus population in the bloodstream reduces to a number less than 0.1. Regarding time in the paper, one time unit means one viral life cycle calculation. In a real life scenario, this could be between half-a-day and a day, depending on the specific viral type [1].

Below we show the evolution viral population with time in the bloodstream and inside cells for the three antiviral strategies discussed in the paper.

### 2 Targeting Fecundity

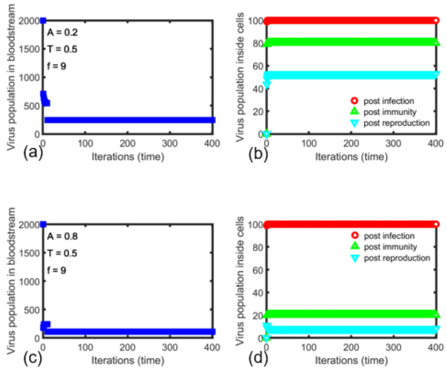


Figure 1: Virus Population in the bloodstream and inside cells for f=9. The therapeutics applied reduces fecundity to 9 after 10th iteration. Top row corresponds to low immunity and bottom row to high immunity.

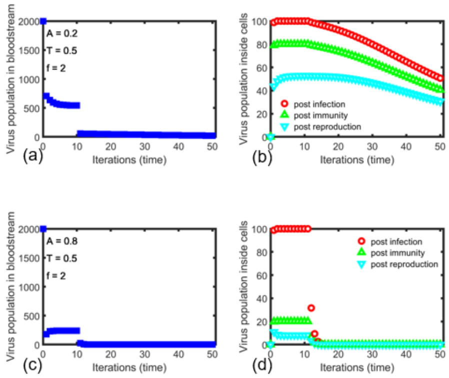


Figure 2: Virus Population in the bloodstream and inside cells. The therapeutics applied reduces fecundity to 2 after 10th iteration. Top row corresponds to low immunity and bottom row to high immunity..

#### 3 Targeting Reproduction Rate , R

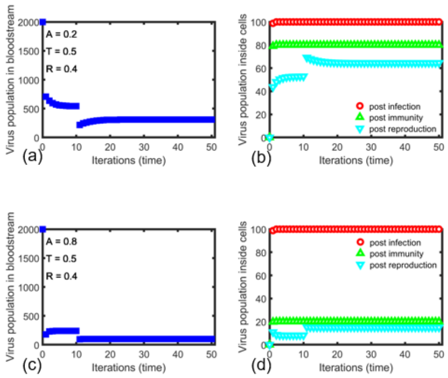


Figure 3: Virus Population in the bloodstream and inside cells. The therapeutics applied reduces reproduction rate R to 0.4 after 10th iteration. Top row corresponds to low immunity and bottom row to high immunity.

### 4 Targeting Infection Rate , I

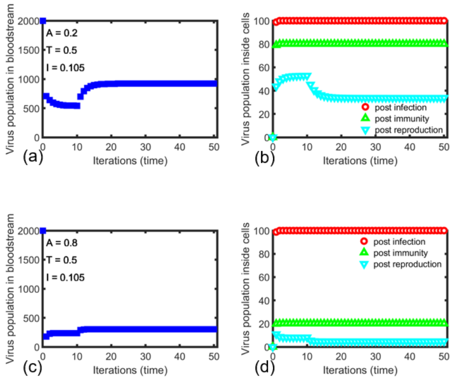


Figure 4: Virus Population in the bloodstream and inside cells. The therapeutics applied reduces infection rate I to 0.105 after 10th iteration. Top row corresponds to low immunity and bottom row to high immunity.

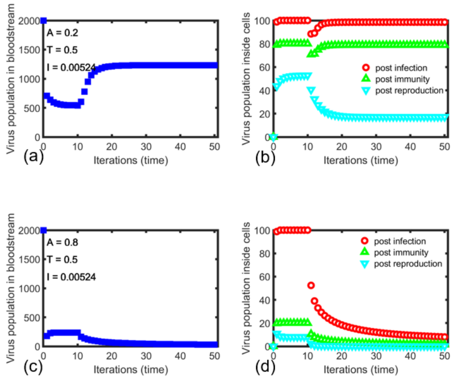


Figure 5: Virus Population in the bloodstream and inside cells. The therapeutics applied reduces infection rate I to 0.00524 after 10th iteration. Top row corresponds to low immunity and bottom row to high immunity.
